# SARS-CoV-2 convergent evolution as a guide to explore adaptive advantage

**DOI:** 10.1101/2021.05.24.445534

**Authors:** Jiří Zahradník, Jaroslav Nunvar, Gideon Schreiber

## Abstract

Much can be learned from 1.2 million sequences of SARS-CoV-2 generated during the last 15 months. Out of the overwhelming number of mutations sampled so far, only few rose to prominence in the viral population. Many of these emerged recently and independently in multiple lineages. Such a textbook example of convergent evolution at the molecular level is not only curiosity but a guide to uncover the basis for adaptive advantage behind these events. Focusing on the extent of the convergent evolution in the spike (S) protein, our report confirms that the most concerning SARS-CoV-2 lineages carry the heaviest burden of convergent S-protein mutations, suggesting their fundamental adaptive advantage. The great majority (21/25) of S-protein sites under convergent evolution tightly cluster in three functional domains; N-terminal domain, receptor-binding domain, and Furin cleavage site. We further show that among the S-protein receptor-binding motif mutations, ACE2 affinity-improving substitutions are favored. While the probed mutation space in the S protein covered all amino-acids reachable by single nucleotide changes, substitutions requiring two nucleotide changes or epistatic mutations of multiple-residues have only recently started to emerge. Unfortunately, despite their convergent emergence and physical association, most of these adaptive mutations and their combinations remain understudied. We aim to promote research of current variants which are currently understudied but may become important in the future.

## Introduction

SARS-CoV-2 virus evolution is shaped by selection imposed by the host and the environment, resulting in new variants with adaptive advantage rapidly taking over previous strains. Global attention is now focused on a number of major variants of concern: B.1.1.7, initially prominent in the United Kingdom, B.1.351, discovered in South Africa, P.1 which has spread rapidly in the State of Amazonas and most recently B.1.617.1/2 spreading from India. Independent acquisitions of S-protein substitutions L452R, E484K/Q, N501Y, and Q677H in these and other lineages were analyzed in great detail (1–5). The intra-host SARS-CoV-2 genomic diversity (6) and the viral evolution in immunocompromised patients (7–10) also received a lot of attention. Yet studies focused on convergent (syn. parallel) evolution highlighting new potential mutations of interest and their combination are missing. Here we analyzed the convergent evolution of SARS-CoV-2 spike protein (S-protein) amino acid (AA) changes which have emerged independently since late 2020 in three or more prominent lineages. In addition, an exhaustive analysis of all possible S-protein receptor-binding motif substitutions, which are reachable by single- and double-nucleotide mutations, was conducted with respect to their global presence and binding effect. Our findings map the peculiarities of the SARS-CoV-2 mutational landscape and reinforce the need of careful monitoring of SARS-CoV-2 evolution.

## Results

To monitor SARS-CoV-2 evolution, we briefly looked for convergent changes among all genes of the SARS-CoV-2 genome at NextStrain (https://nextstrain.org/ncov/global) (5). We identified convergent evolution in S-protein, ORF1a, ORF1b, ORF3a, M, ORF8, N, and ORF9b. Since all of the most concerning variants contain multiple mutations in the S-protein, and since all globally used vaccines *(i.e.* mRNA- and adenovirus-based) immunize exclusively against this protein, we further analyzed its amino acid changes, as recorded in the GISAID database (11). The total number of mutations, including indels, at a given position, was calculated and plotted (Figure 1). This identified a median number of 80 sequences of mutated AA per position, with only three percent of AAs out of the total 6,554,383 substitutions detected in more than 10^4^ genomic sequences (out of 1.2 mil. genomic sequences in GISAID; May 5, 2021). Enrichment of mutations at a given position can result from fixation of a variant which quickly becomes dominant, as for example D614G, whose positive impact on virus fitness was demonstrated (12); this is particularly common for mutations at early stages of a pandemic. A second cause of a rise of certain mutations to high prevalence is their convergent evolution, where the same mutations arise independently in different viral lineages. The selection of these mutations often accompanies the emergence of novel, successful variants.

**Figure 1.**
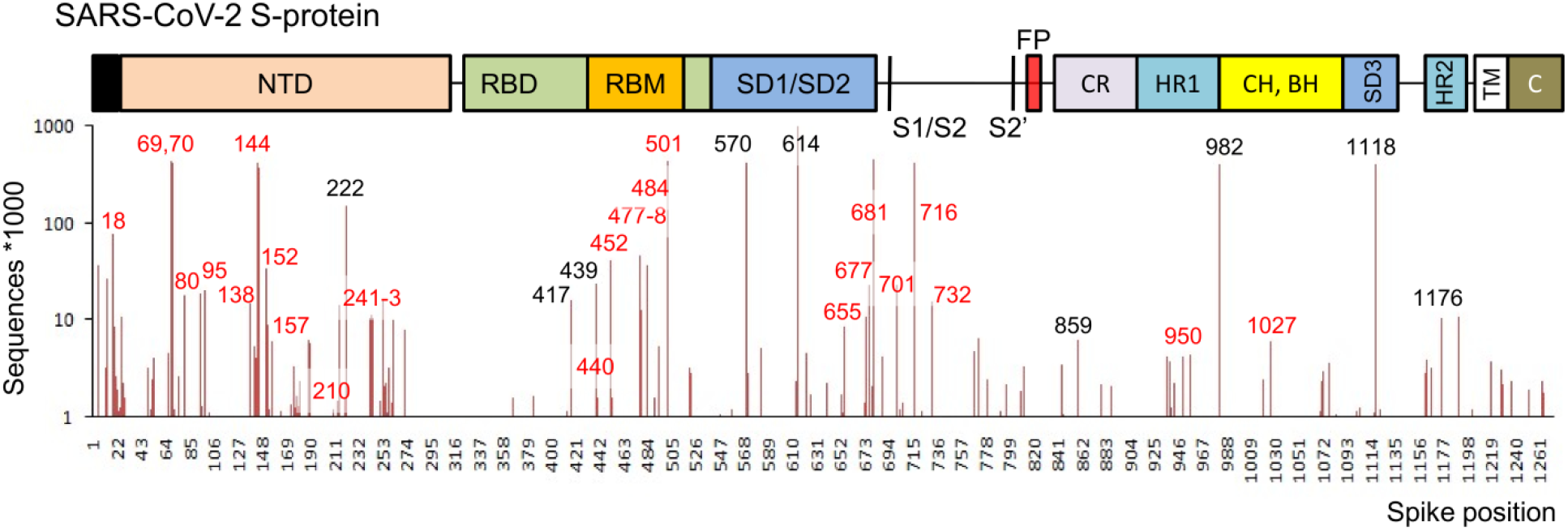
SARS-CoV-2 Spike protein domain organization and acquired mutations. Domain organization (upper part): NTD – N-terminal domain, RBD – receptor-binding domain, RBM – receptor-binding motif, SD – subdomain, S1/S2/S2’-cleavage sites, FP – fusion peptide, CR – connecting region, HR – heptad repeat, CH – central helix, BH – β-hairpin, TM – transmembrane helix, C – cytoplasmic tail; The numbers are given for prevalent mutations. In red are denoted positions with convergent evolution (≥3 distinct lineages) and black numbers for those without convergent evolution. Lines show mutations sequenced > 1000 (May 5, 2021).

A comparative analysis was conducted to unravel the approximate extent of convergent evolution in individual S-protein AAs, identifying 40 mutations at 25 sites which have emerged independently at least 3-times in unrelated viral lineages (Table 1; for details, see Methods.). These mutations are located in three distinct hotspots – N-terminal domain (NTD), receptor-binding domain (RBD), and Furin cleavage site (Figure 2A). Mutation E484K was present in the highest number of unrelated lineages [11, Table 1], followed by L452R [10], P681H [9], Y144Δ [8], HV69-70Δ [6], Q677H [6] and N501Y [6]; these mutations thus display the strongest convergent evolution. The overall load of convergent mutations in the S-protein was quantified (lineage parallelism score, Table 1). Importantly, the lineages raising the greatest concern also displayed the highest levels of parallelism, reinforcing the proposed adaptive importance of convergent mutations with regard to virus fitness.

**Figure 2.**
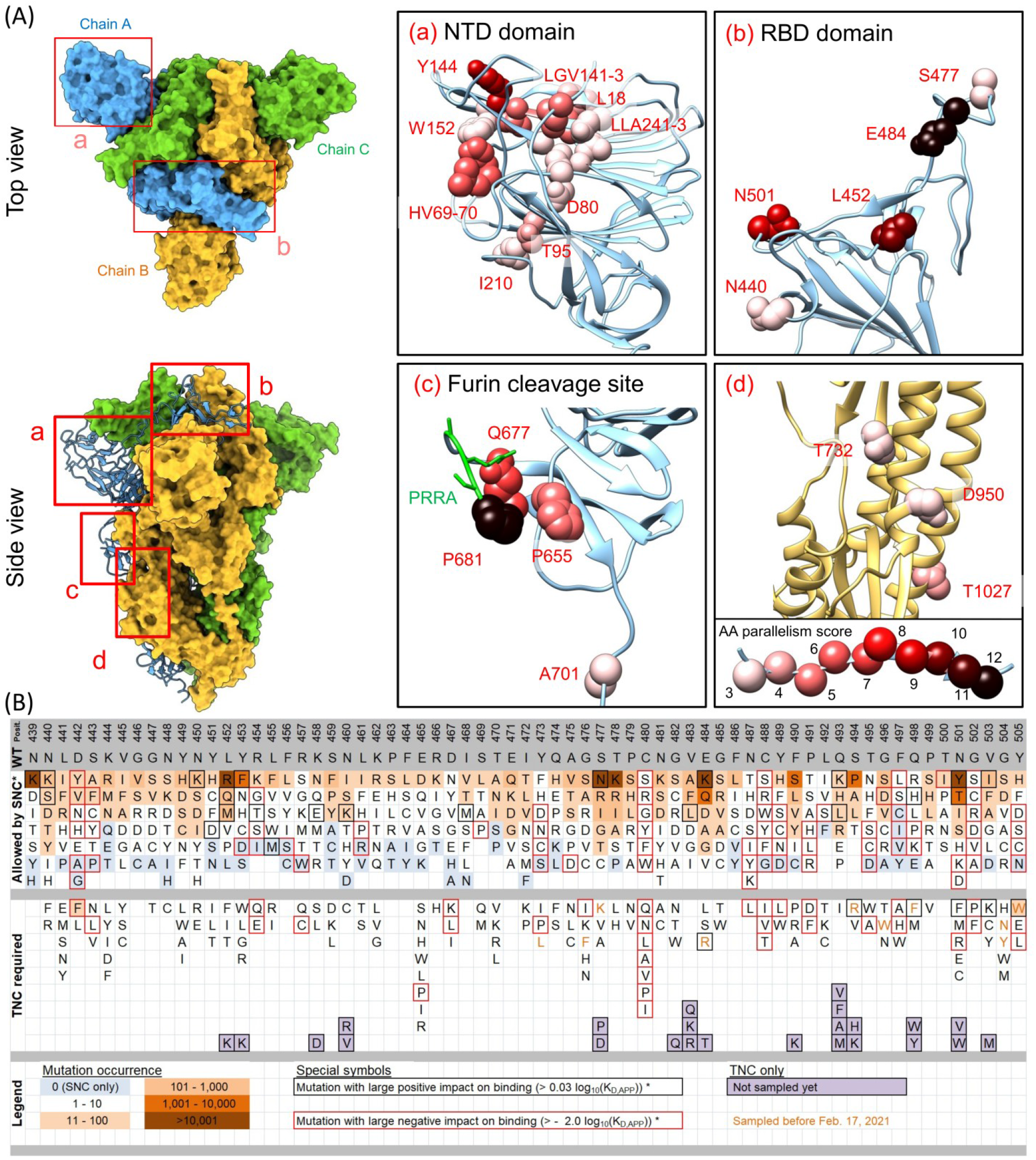
Convergent evolution and mutations in SARS-CoV-2 S-protein. (A) Localization of convergent mutations on the structure of Spike protein (PDB ID: 6zge) and their corresponding parallelism score (for more details see Table 1 and Methods). Green residues in inset c) highlights the Furin cleavage site. Inset d) shows the region covering a portion of heptad repeat, central helix and β-hairpin domains. (B) The mutational sequence space of RBM and its coverage by identified mutations. All possible SNC (single-nucleotide change) AA substitutions are denoted. For TNC (two-nucleotide change) substitutions, only the subset of AA substitutions which were sampled in GISAID are denoted (in background color scale according to the legend), together with substitutions with positive binding impact not yet sampled (violet background)

**Table 1.**
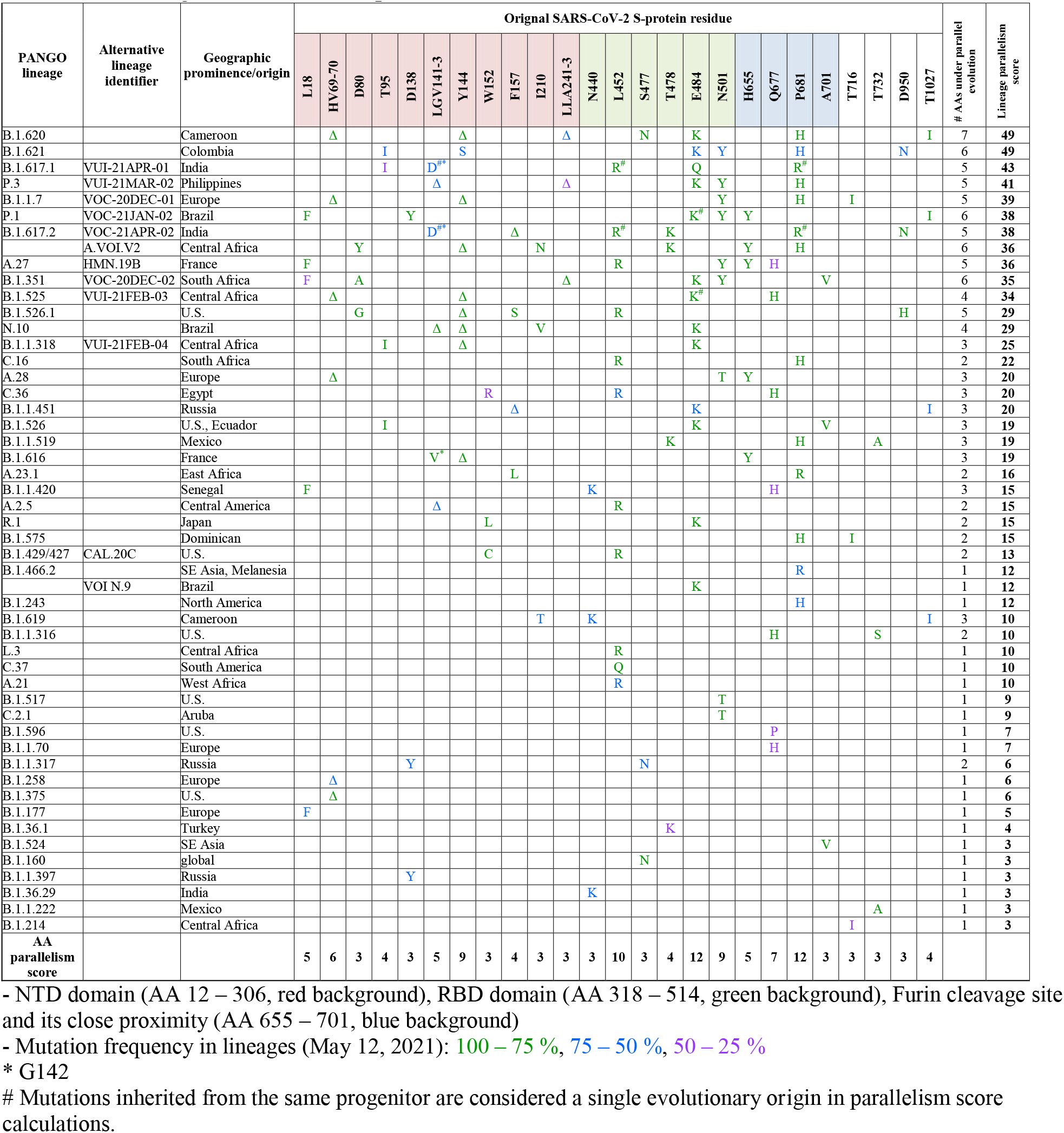
Convergent evolution in S-protein.

Next, we focused on the interaction interface of RBD, the so called receptor binding motif (RBM, AA 439 – 505), to analyze the depth reached by SARS-CoV-2 in exploiting the mutational space. For this, we generated *in silico* all the mutations which can be achieved with a single nucleotide change (SNC) of the original codons. The likelihood of a successive double nucleotide change acquisition in a single codon is much lower, as it requires the single nucleotide as well as the double-nucleotide mutation to be of evolutionary advantage for the lowest frustration trajectory to be sampled. The results (Figure 2B) demonstrate that nearly all possible variants allowed by SNC were already sampled in sequenced SARS-CoV-2 genomes. The alternative residues reachable by two nucleotide changes (TNC) appear among sampled genomes at very minute numbers (<10 total occurrences), yet their repertoire is growing (Figure 2B). Similar to double-nucleotide mutations, epistatic mutations are rarely found among SARS-CoV-2 mutations (13). Most of the AA changes exhibiting positive effect on ACE2 receptor binding [Log_10_(K_D,app_) > 0.03], calculated from deep-mutational scanning (14) (Figure 2B, framed in black) are present at significant frequencies among GISAD genomes (13), reinforcing improved binding as an importing contributor to adaptive fitness gain. Conversely, the selection of most mutations which negatively affect receptor binding (Figure 2B, framed in red) is disfavoured in SARS-CoV-2 evolution.

## Discussion

COVID-19 is the first pandemic disease where unprecedented amount of infectious agent sequence data has been accumulating in real time. This represents a valuable resource to learn about the changes of SARS-CoV-2 as the virus spreads and evolves. Our analysis shows that among the vast number of mutations which have been detected in SARS-CoV-2 viral genomes, only few rose to high frequencies. Strikingly, many of these display convergent evolution, which implies a strong adaptive advantage conferred by the particular mutations. Upon a closer look at the crucial S-protein, three hotspots bearing multiple convergently evolved mutations were identified – NTD domain, RBD domain and the Furin cleavage site (Figure 2A) creating a conspicuous pattern reflecting their putative fitness advantage. Based on the domain location of such mutations we can speculate about their importance. NTD domain plays a role of a “wedge” in controlling the S-protein conformations (15), its interaction with host surface sialosides was suggested (16) and recently the binding to alternative entry receptors were proposed (17). Interestingly, all NTD AAs which display convergent evolution form a perfectly continuous patch, which further stresses the need of their functional analysis. The function of Furin cleavage site is well established (18, 19) as well as the relevance of the convergent Q677H mutation (5). The impact of additional mutations in this region remains to be analyzed but their increased fitness effect due to faster processing can be expected. As with NTD, the Furin cleavage site AAs under convergent evolution are in a direct contact with each other.

Apart from delineating the S-protein regions key for adaptation to the human host, the clustering of AAs under convergent evolution suggests the potential synergic effects between them, as was already observed for the couple E484K/N501Y (13). These mutations should thus be evaluated in their combinations, due to their increasingly observed cooccurrence in individual lineages *(e.g.* B.1.1.7 with E484K designed as VOC-202102/02). Our report should help to identify the worrisome novel lineages hosting combinations of S-protein mutations under positive selection and thus prevent their neglected spread, as happened for some recent lineages (*e.g.* B.1.620)(20). Structural biology studies, the golden standard for bioinformatics-based functional predictions, show a strong bias towards the most attractive mutations. Among the 355 structures of SARS-CoV-2 Spike protein (to 4.5.2021, PDBe-KB COVID-19 Data Portal), we identified only D80A (representative PDB ID: 7lyn (21)), K417N/T (7lym (21)), S477N (7bh9 (13)), E484K(7lww (21)), N501Y(7mjg (2)) and T716I (7lwu (21)) mutations, which together represent only a small fraction of residues which need to be analyzed in further details.

Strikingly, most of the affinity-enhancing mutations (Figure 2B, black-framed) reachable by SNC are already abundant in the global genomic dataset. In contrast, we identified only a single mutation (Y505W) with a relatively high representation and affinity-enhancing performance compared to the wild-type, which requires two nucleotide changes. Many TNC mutations with apparently tighter binding to ACE2 have not yet been sampled (Figure 2B, violet background). The comparison with TNC sampled by 17.2.2021 (Figure 2B, residues in orange) shows a dramatic increase in the total wealth of these advanced mutations during the last three months, albeit still with very low frequencies. This also explains why epistatic mutations, which require orchestrated changes in a number of nucleotides in the protein are still rare. In contrast, SNC mutations which improve binding and have so far been sampled in zero or negligible frequencies are likely restricted by structural constrains. The deep-mutational scanning (14) suggests that a significant proportion of non-sampled or low-occurrence SNC mutations substantially decrease binding (Figure 2B, red-framed). Overall, these observations are in line with our previous work, suggesting that receptor binding and the acquisition of positive charge on the RBM have been until now the strongest driver for positive selection (13). The mutations that do not follow this trend are L452R, F490S, T478K, and S494P; we thus speculate that their benefits for viral fitness are likely connected with the host immune system. The immunocompromised patients, treated or non-treated with convalescent plasma or neutralizing antibodies, have been regarded crystal balls for prediction of viral evolution (7, 8), showing the emergence of N501Y, S494P, Q493K, Y489H, E484K, T478K and N440D (9, 10) variants during infection. Surprisingly, only mutations E484K and N501Y arose convergently among patients, and their emergence was restricted to a low number of patients. Furthermore, no combination of 484 and 501 was observed only in a single patient, suggesting a stepwise selection of multiple residues possibly linked with transmission benefit. On the other hand, the deletions Δ141-143, which was observed multiple times in different patients (10, 22) and_ ΔY144 (23) were clearly selected in the response to host environment. It is evident that every mutation contributes by multiple different parameters and mechanisms which need to be evaluated systematically.

So far, the evolutionary pressure on SARS-CoV-2 was mainly to increase virus fitness in a virgin environment. However, the global introduction of vaccines is expected to shift the pressure towards immune-evasion mutation, even on the expense of fitness. The predictability of convergent S-protein evolution could in principle increase our odds that the S-protein sequences utilized in the universal second-generation vaccines, when carefully selected, will effectively protect the global population from the landscape of viral variants of current and future concern.

## Materials and Methods

### Convergent evolution analysis

We collected all S-protein AA mutations present in SARS-CoV-2 lineages of concern which were described to have emerged between autumn 2020 (*e.g.* B.1.1.7) and spring 2021 (*e.g.* B.1.621). The distribution of each mutation was visually examined in the representative global subsample of approximately 4,000 SARS-CoV-2 genomes (https://nextstrain.org/ncov/global) (24). Through AAs which were mutated in at least three genetically distinct lineages, further lineages were collated and all their S-protein AA mutations were subjected to the same procedure. After several iterations, a finite set of 50 lineages carrying convergent S-protein AA mutations was established. The frequency of S-protein AA mutations among genomes present in GISAID (11) was determined using the Lineage Comparison tool (https://outbreak.info/)(25). The AA parallelism score was established for all convergently mutated AAs, by summing their independent emergence events (≥25 % within-lineage frequency). The lineage parallelism score was calculated by summing the parallelism scores of S-protein AAs mutated in individual lineages.

### Spike RBM mutations exhaustive analysis

All amino acid substitutions and occurrences in the S-protein were downloaded from the GISAID database (5 May 2021). In addition, the dataset for RBM was downloaded on 17^th^ February 2021. The data manipulations as well as all SNC changes generation were done by in-house written Python 3.7 code. Deep-mutational scanning ΔLog_10_(K_D, App_) values were extracted from https://jbloomlab.github.io/SARS-CoV-2-RBD_DMS/ (14). 3D visualization and analyses were performed using UCSF Chimera 1.15 (26) or ChimeraX 1.2 (27). The missing structures in PDB 6zge were modeled by Modeller suite implemented in Chimera (28).

## Author Contributions

Author contributions: J.Z. and G.S. conceived the project; J.Z., J.N. and G.S. performed experiments; J.Z., J.N. and G.S. wrote the manuscript.

## Acknowledgments

Funding: This research was supported by the Israel Science Foundation (grant No. 3814/19) within the KillCorona – Curbing Coronavirus Research Program and by the Ben B. and Joyce E. Eisenberg Foundation. J.N. acknowledges support by project MICOBION (H2020 No 81022), funded by Research Executive Agency (REA), and by the European Regional Development Fund and the Ministry of Education, Youth and Sports of the Czech Republic, grant number CZ.02.1.01/0.0/0.0/16_019/0000785.

